# Ordered deployment of distinct ciliary beating machines in growing axonemes of vertebrate multiciliated cells

**DOI:** 10.1101/2022.07.05.498879

**Authors:** Chanjae Lee, Yun Ma, Fan Tu, John B. Wallingford

## Abstract

The beating of motile cilia requires to coordinated action of diverse machineries that include not only the axonemal dynein arms, but also the central apparatus, the radial spokes, and the microtubule inner proteins. These machines exhibit complex radial and proximodistal patterns in mature axonemes, but little is known about the interplay between them during motile ciliogenesis. Here, we describe and quantify the relative rates of axonemal deployment for these diverse cilia beating machineries during the final stages of differentiation of *Xenopus* epidermal multiciliated cells.

## Introduction

The coordinated beating of motile cilia on multiciliated cells (MCCs) drives directional fluid flow that is essential for the development and homeostasis of diverse organ systems, notably the vertebrate airways, reproductive tract, and central nervous system (Brooks and Wallingford, 2014; Mahjoub et al., 2022). Beating of these cells’ cilia relies on several protein machines that are radially organized throughout the axoneme (Fig. 1, left)(Dutcher, 2020). For example, Microtubule Inner Proteins (MIPs) project inside the outer doublet microtubules, provide structural stability, and likely regulate axonemal dynein function (Ichikawa and Bui, 2018). The axonemal dyneins, in turn, reside outside these doublet microtubules, and include both the outer and inner dynein arms (ODAs, IDAs)(King, 2018; Yamamoto et al., 2021). In the center of the axoneme are the central pair microtubules and these are surrounded by the central apparatus (Loreng and Smith, 2017). Finally, radial spokes project from the doublets toward the central pair (Zhu et al., 2017). Each of these systems are required for normal ciliary beating, but because each system is more frequently studied in isolation, the interplay between them remains ill defined.

**Figure 1:**
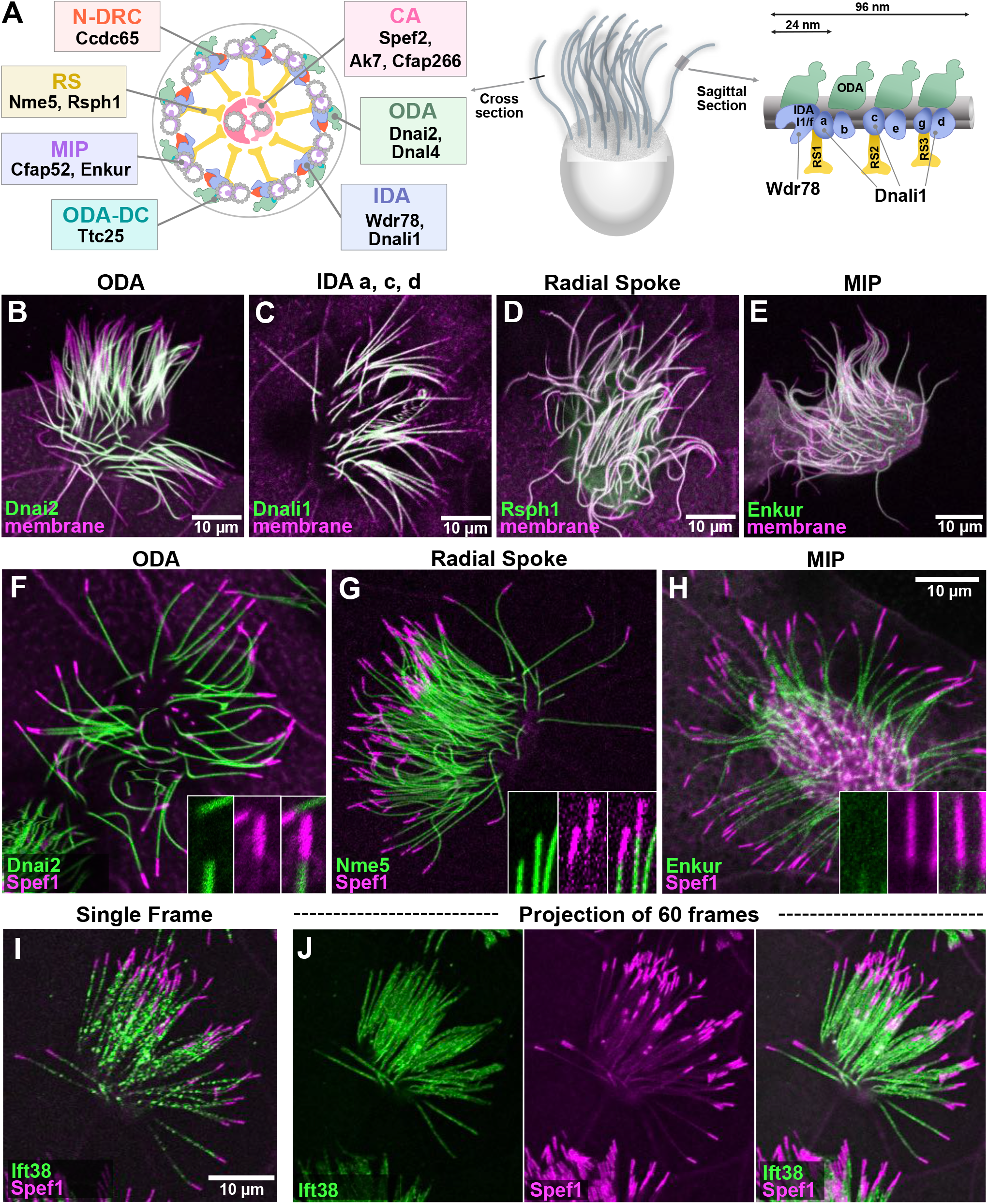
A) Schematics of an MCC axoneme, with relevant components of ciliary beating machineries indicated. ODA (outer dynein arm), IDA (inner dynein arm), CA (central apparatus), N-DRC (nexin-dynein regulatory complex), RS (radial spoke), MIP (microtubule inner protein), ODA-DC (ODA docking complex) B-E) The indicated ODAs, IDAs, radial spokes, and MIPs do not extend to the distal limit of the axoneme, as marked by membrane RFP. F-H) The indicated ODA, radial spoke, and MIP are excluded from the distal Spef1-enriched domain. I) A single frame from a time-lapse movie of GFP-Ift38 and RFP-Spef1. J). A projection of 60 frames of time-lapse data reveals absence of Ift38 from the Spef1-enrched distal domain.

In addition to the radial patterning, motile axonemes display a complex proximal to distal (PD) patterning. Such patterning is most obviously displayed by the presence of ODAs and IDAs in highly order repeating units (Fig. 1A, right), but there are more complex patterns as well (Dutcher, 2020). For example, the PD patterning of specific sub-types ODAs and IDAs has been well defined by structural studies in *Chlamydomonas* (Bui et al., 2012; Piperno and Ramanis, 1991; Yagi et al., 2009) and by immunostaining of ODA/IDA heavy chains in vertebrates (Dougherty et al., 2016; Fliegauf et al., 2005; Oltean et al., 2018). Motile cilia, including those on vertebrate MCCs, also display specialized distal tip domains that are distinct in structure and composition from the rest of the axoneme (Soares et al., 2019).

One of many unanswered questions relates to how these radial and proximodistal patterns are established as new cilia grow during the development of MCCs. For example, a small subset of proximally-restricted IDAs are known to be deployed only very late during cilia regeneration in *Chlamydomonas* (Piperno and Ramanis, 1991) and immunostaining has identified certain distinct heavy chains associated with these motors (Yagi et al., 2009). A recent study has also explored the initial deployment of certain ODA and IDA heavy chains during ciliary growth in human MCCs (Oltean et al., 2018). But little or nothing is known about the timing of deployment of ODA and IDA light or intermediate chains or of how ODA/IDA deployment relates to that of distinct machineries like the radial spokes of central apparatus. Likewise one paper has carefully defined central apparatus assembly during flagellogenesis in *Chlamydomonas* (Lechtreck et al., 2013), but no comparable study has been made in MCCs.

These questions are important, because recent studies have highlighted the functional interplay between the central apparatus and deployment of ODA/IDAs and radial spokes (Konjikusic et al., 2022) and because disruption of cytosolic assembly factors for axonemal dyneins in human motile ciliopathy elicits proximodistally-restricted defects in ODA/IDA deployment (Lee et al., 2020; Omran et al., 2008; Tarkar et al., 2013; Yamaguchi et al., 2018). As a starting point for future experimental studies, we present here a quantitative description of the relative rates of deployment of MIPs, radial spokes, the central apparatus, ODAs, and IDAs in the growing axonemes of developing MCCs in the *Xenopus* epidermis.

## Results

### Cilia beating machines and IFT are excluded from the distal axoneme of *Xenopus* MCCs

The *Xenopus* epidermis provides a tractable vertebrate model for imaging MCC development (Brooks and Wallingford, 2015; Walentek and Quigley, 2017; Werner and Mitchell, 2012). As a starting point, we co-expressed GFP fusions to ciliary beating machines together with membrane-RFP (memRFP) to mark the axoneme. Using full-length, mature axonemes at St. 27. We first examined representative markers of the ODAs and IDAs (Dnai2, Dnali1)(King, 2018), the radial spokes (Rsph1)(Anderegg et al., 2019), and MIPs (Enkur)(Gui et al., 2021; Khalifa et al., 2020). All of these reporters displayed an obvious lack of signal in the distalmost ∼10% of the axoneme (Fig. 1B-E).

In organisms from *Tetrahymena* to humans, motile cilia display specialized distal tip regions (Soares et al., 2019), and in *Xenopus* epidermal MCCs this regions is reliably marked by a strong enrichment of Spef1 (aka Clamp1)(Gray et al., 2009). This microtubule bundling protein is also present in the central apparatus but at much lower intensity, and its distally enriched domain is readily identifiable (Konjikusic et al., 2022). To ask if the distal region of MCC axonemes lacking ciliary beating machinery corresponds to the Spef1-enriched domain, we co-expressed Spef1-RFP with GFP fusions to other ciliary proteins. In mature axonemes, we found that the ODA subunit Dnai2, the radial spoke protein Nme5, and the MIP Enkur each displayed the same pattern, with the distal GFP signal terminating at the proximal limit of the enriched Spef1-RFP signal (Fig. 1F-H).

Dynein arms, radial spokes, and other cargoes are actively transported within axonemes by the IntraFlagellar Transport (IFT) system (Lechtreck, 2015), so we asked how the movement of IFT trains related to the Spef1-enriched domain. We used high speed time-lapse imaging to track the movement of the IFT protein, Ift38/Cluap1 (Fig. 1I) (Lee et al., 2014), and by merging several frames of such movies, we clearly observed that the distal limit of IFT movement matched tightly to the proximal limit of the Spef1-enriched domain (Fig. 1J).

These data lead us to several interesting conclusions: First, all four major elements of ciliary beating machinery display similar distal extents in mature axonemes. Second, consistent with the findings in *Chlamydomonas* that IFT is the key mechanism for transport of dynein arms and radial spokes within cilia (Lechtreck, 2015), the distal extent of IFT is matched to that of the ciliary beating machinery.

Third, the Spef1-enriched region in mature axonemes represents a specialized distal domain that lacks components of the ciliary beating machinery as well as the IFT machinery that transports them. Finally, the absence of IFT from the distal Spef1-enriched domain is consistent with the finding that Eb1 is also localized in this distal region in *Xenopus* (Konjikusic et al., 2022) and the same distal enrichment of Eb1 in *Chlamydomonas* is independent of IFT (Harris et al., 2015).

### The specialized distal axoneme forms early during axoneme growth and is not coupled to deployment of the ciliary beating machinery

*Xenopus* epidermal MCCs arise via radial intercalation at the early tailbud stage and immediately begin to grow cilia, and over the next several hours, immature cilia can be readily observed (Drysdale and Elinson, 1992; Stubbs et al., 2006). To ask how the Spef1-enriched distal domain was assembled during ciliary growth, we examined RFP-Spef1 and memGFP during cilia growth (St. 21-27). At the earliest stages examined when cilia were quite short, Spef1 weakly labeled axonemes along their length, but by st. 22, a small region of distal enrichment was visible (Fig. 2A, B, F). This distal enrichment grew as axonemes grew, finally reaching it’s final size by st. 27 (Fig. 2C, F). To understand how the assembly of the Spef1-enriched domain relates to axonemal deployment of ciliary beating machinery, we co-expressed RFP-Spef1 and the radial spoke protein, Nme5-GFP (Anderegg et al., 2019) and imaged growing cilia.

**Figure 2:**
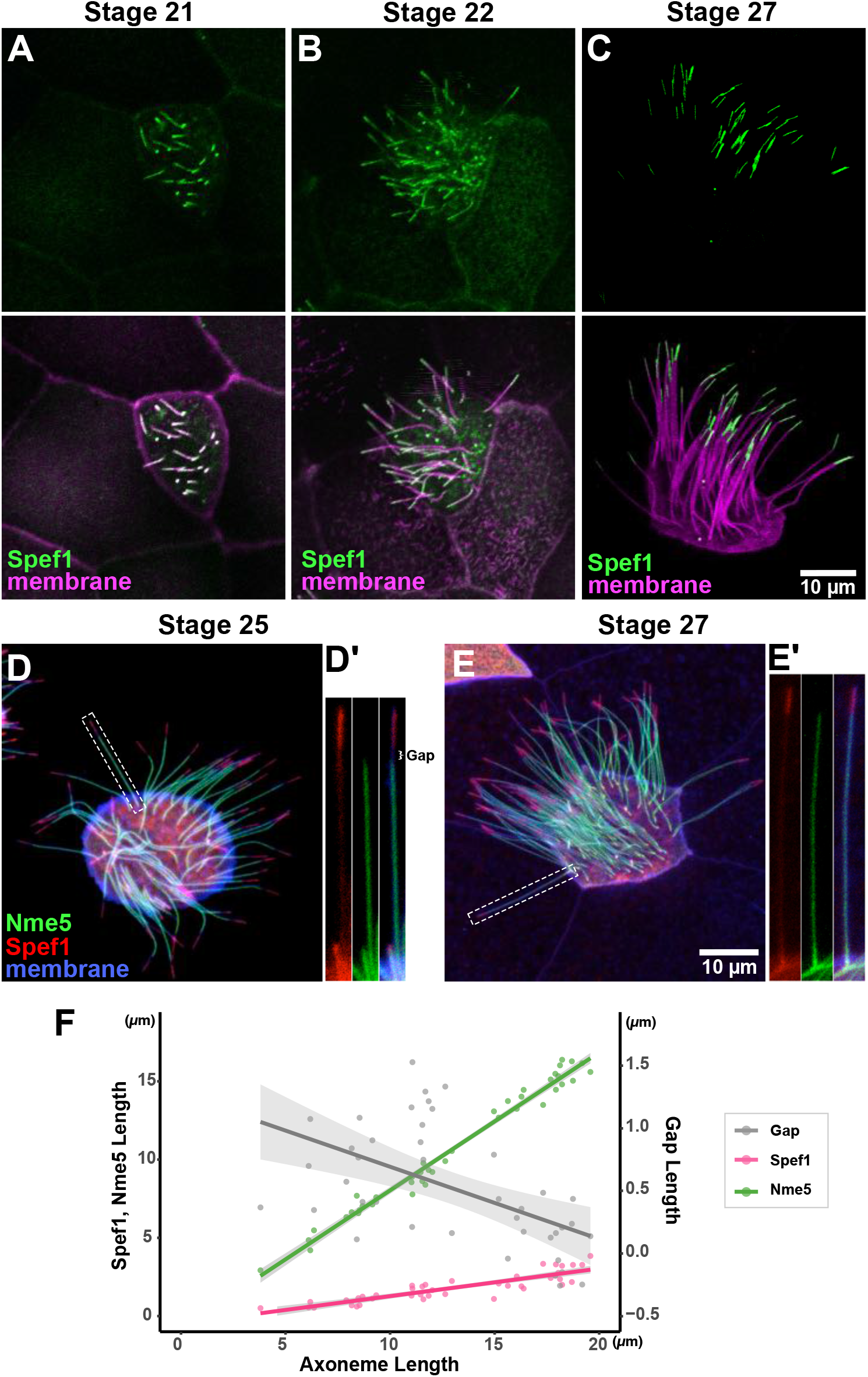
A-C) Spef1-GFP distal enrichment in growing axonemes at stages indicated. D-E) The deployment of the radial spoke protein Nme5 does not reach to the Spef1-enriched domain at st. 25 but does so by st. 27. F) Plot of axoneme length versus Spef1 length, Nme5 length and the length of the gap separating them.

Interestingly, in shorter growing cilia we observed a clear gap between the distal limit of Nme5 and the proximal limit of Spef1 (Fig. 2D, inset at right). As cilia approach their final length, this gap shrank and eventually disappeared so that Nme5 abuts the Spef1-enriched domain in full-length, mature cilia (Fig. 2E, F). These data suggest that as the axoneme grows, distal tip assembly and deployment of the beating machinery are uncoupled, with the distal Spef1-enriched domain assembling from the growing tip of the axoneme and the relatively delayed deployment of radial spokes proceeding from the ciliary base before finally terminating at the proximal end of the Spef1-enriched region.

### Deployment of microtubule inner proteins and radial spokes and is tightly coupled to axoneme elongation, while the central apparatus lags behind

We next sought to determine the timing of deployment of ciliary beating machines into the axoneme during ciliary growth. While methods for physically immobilizing *Xenopus* MCC cilia are well established (Brooks and Wallingford, 2015; Werner and Mitchell, 2013), we have not found them to be effective for time lapse analysis of individual growing cilia. We therefore developed a new approach for quantifying the *relative* rates of ciliary beating machinery deployment in growing axonemes.

We used GFP fusions to visualize ciliary proteins and co-expressed memRFP to visualize the entire axoneme. Because all elements of the beating machinery ultimately extend to similar distal positions in *mature* axonemes (Fig. 1), relative rates of deployment can be assessed by examining *growing* axonemes of differing lengths and plotting the coverage of GFP-decoration against total axoneme length. Since mature axonemes on *Xenopus* MCCs are ∼15∼20 microns in length, we considered only those <8 microns in length here to ensure that we analyzed only actively growing axonemes.

We used this method first to examine two MIPs, Enkur and Cfap52 (Dougherty et al., 2020; Sigg et al., 2017), and we found that each displayed essentially a one-to-one correlation to overall length during axoneme growth (Fig. 3A). This result is consistent with the fact that Enkur and Cfap52 are tightly embedded within the outer microtubule doublets of the axoneme (Gui et al., 2021; Li et al., 2022). The radial spoke protein Nme5 displayed similarly swift kinetics, extending at the same rate as the MIPs and the overall axoneme (Fig. 3B).

**Figure 3:**
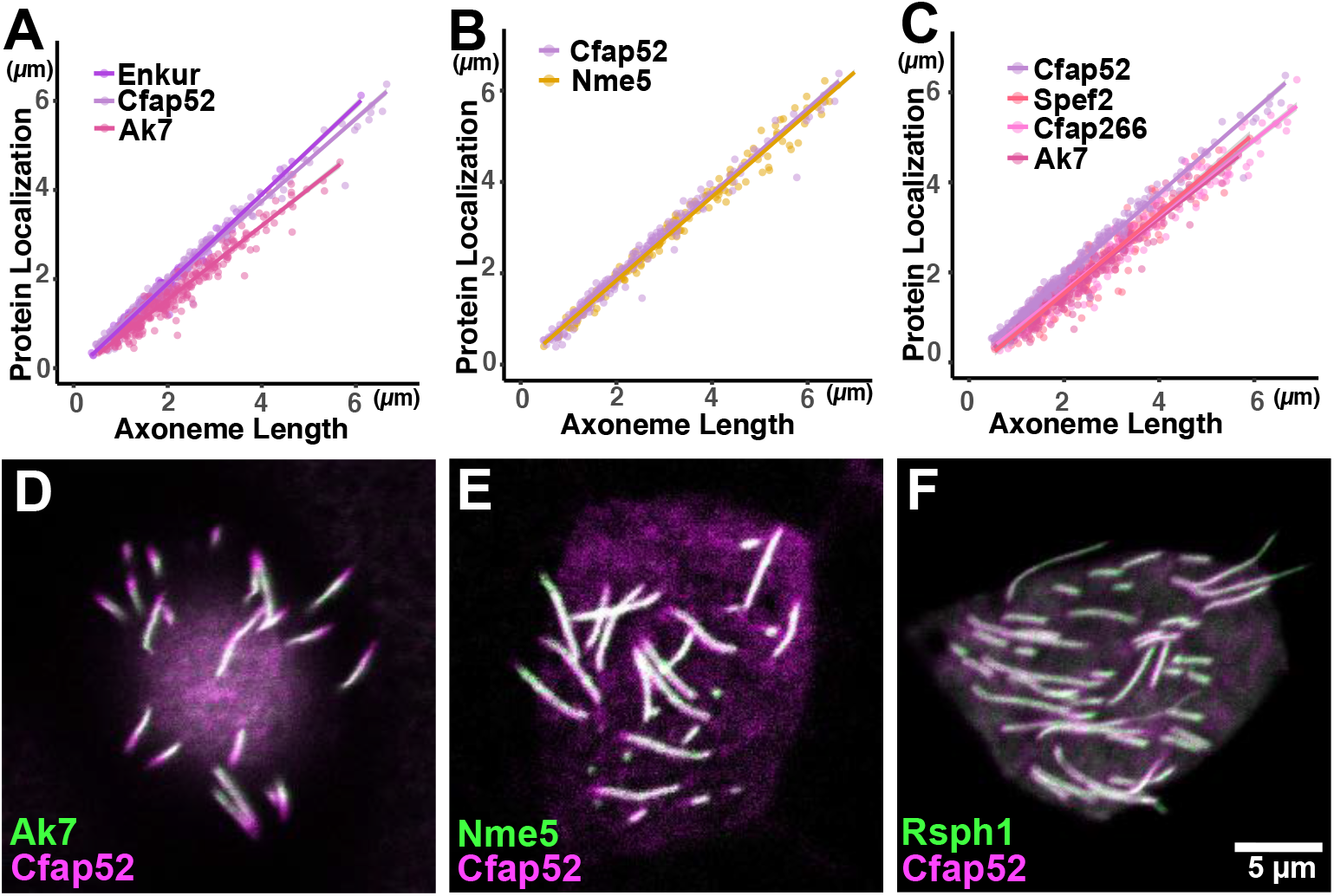
A-C) Plots of total axoneme length versus the length of the axoneme decorated by the labeled protein. D) Double labeling shows that Cfap52 (magenta) consistent extends distally to Ak7 in immature axonemes, consistent with their distinct kinetics in A. E, F). Nme5 and Rsph1 do not display consistent differences from Cfap52, consistent with their similar kinetics in B-C).

By contrast, the central apparatus protein Ak7 displayed distinct kinetics, lagging well behind Enkur and Cfap52 (Fig. 3A). The central apparatus is subdivided into C1 and C2 regions (Loreng and Smith, 2017), and Ak7 is a component of C2 region. It’s notable then that another C2 protein, Cfap266, and a C1 protein, Spef2 (Dai et al., 2020; Zhao et al., 2019) also displayed identical kinetics (Fig. 3C). (Note: To facilitate comparisons, certain data are re-plotted in multiple panels. Summary statistics for all data are presented in Fig. 5, below).

To confirm these relative rates of deployment in axonemes, we co-expressed RFP and GFP fusions to representative pairs of ciliary proteins. For example, we consistently observed a Cfap52-RPF-*positive* and Ak7-GFP-*negative* region in the distal axoneme of cells co-expressing the two (Fig. 3D), consistent with Ak7 deployment lagging that of Cfap52 (Fig. 3A). By contrast, when we co-expressed Nme5-GFP and Cfap52-RFP, we observed no consistent extension of one reporter distal to the other (though RFP-only or GFP-only regions were sometimes observed (Fig. 3E). Another radial spoke protein, Rsph1 displayed a similar pattern (Fig. 3F). Thus, the kinetics of MIPs and radial spokes suggest that they are directly integrated into the growing axoneme as it is assembled, while the central apparatus is delayed in its deployment.

### Deployment of dynein arms is delayed relative to axoneme elongation, and inner and outer motors display distinct kinetics

We next examined the deployment of dynein arms by co-expressing ODA or IDA subunits fused to GFP together with memRFP (Fig. 4A, B) and observed interesting differences in their kinetics. First, deployment of the ODA intermediate chain Dnai1 lagged well behind Enkur and overall axoneme growth as reported by memRFP (Fig. 4C), and when co-expressed, the Enkur-GFP signal consistently extended more distally than did Dnai2-GFP (Fig. 4D). Another ODA subunit, the light chain Dnal4, displayed kinetics identical to Dnai2 (Fig. 4C), together suggesting that ODA deployment lags well behind axoneme elongation.

**Figure 4:**
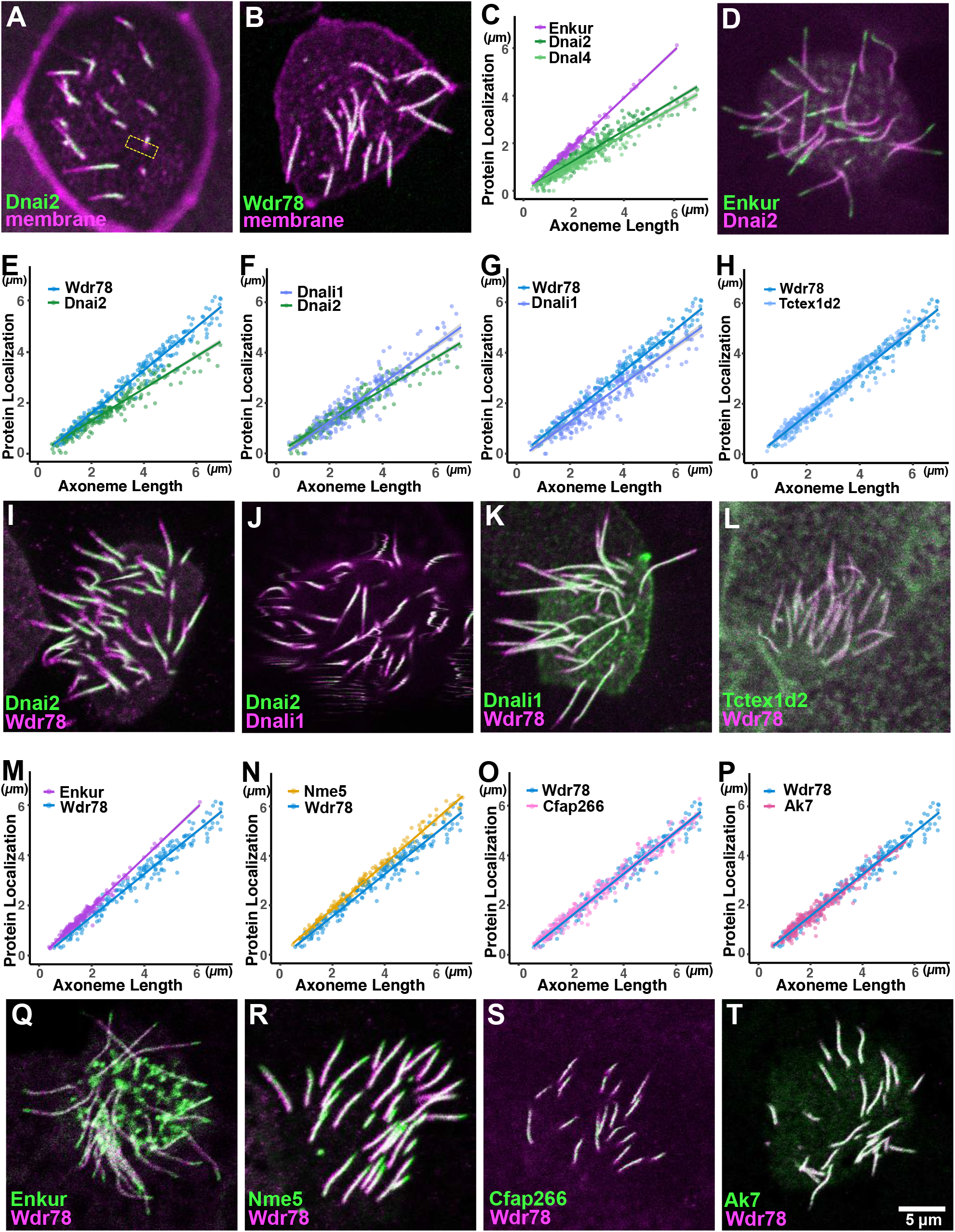
A, B) Dnai2 and Wdr78 are clearly absent from the distal region of immature axonemes. C) ODA subunits display far slower deployment kinetics as compared to the MIP, Enkur. D) Dual labeling reveals that Enkur consistently extends distally to Dnai2 in growing axonemes, consistent with their distinct kinetics in C. E-H) IDAs display faster deployment kinetics than ODAs, and IDA sub-types display distinct kinetics. I-L) Dual labeling confirms relative rates of deployment for ODA/IDA subunits. M-P) IDAs deploy at rates slower than MIPs and radial spokes, and similar to the central apparatus. Q-T) Dual labeling confirms kinetic data on rates of IDA deployment.

While ODAs drive ciliary beating, a larger and more diverse set of IDAs tune the waveform (Fig. 1A, right)(Yamamoto et al., 2021). We next examined the IDA(f) subunit Wdr78 and the IDA(a,c,d) subunit Dnali1 and found that each deployed substantially faster than did the ODAs (Fig. 4E, F), though the two IDAs were not identical. Indeed, Dnali1 lagged subtly behind Wdr78 (Fig. 4G). Another IDA(f) subunit, Tctex1d2 displayed kinetics identical to Wdr78 (Fig. 4H), suggesting the distinct kinetics between Dnali1 and Wdr78 does not related to specific protein subunits but rather more generally to IDA sub-types. Each of these results was confirmed by dual-labeling (Fig. 4I-L).

For a more comprehensive comparison, we then plotted the kinetics of IDA(f) subunit Wdr78 relative to the MIP Enkur and radial spoke protein Nme5 and found that Wdr78 lagged subtly behind (Fig. 4M, N). This result prompted us to compare IDA(f) to the central apparatus, and we found that Wdr78 displayed kinetics identical to those displayed by Cfap266 or Ak7 (Fig. 4O, P). These results, too, were confirmed by dual-labeling experiments (Fig. 4Q-T).

This series of results raises several interesting points: First, deployment of all IDA and ODA subunits examined here lagged behind Enkur, and thus behind the axoneme generally, suggesting that neither IDA nor ODA deployment is directly integrated with axoneme growth. Second, the kinetics for two ODA subunits were similar, as were those for two IDA(f) subunits, consistent with the fact that axonemal dyneins are pre-assembled in the cytoplasm and deployed to axonemes as intact motors (Qiu and Roy, 2022). Third, the divergent kinetics of ODA, IDA(f), and IDA(a,c,d) suggest those intact motors are deployed to the axoneme independently.

### Divergent patterns of deployment kinetics for other elements of the beating machineries

Thus far, a certain coherence can be perceived in the relative rates of deployment for ciliary beating machines, as multiple MIPs share kinetics, as do multiple central apparatus proteins, IDA(f) subunits or ODA subunits). However, this coherence did not extend to additional elements of the cilia beating machinery. For example, the Nexin Dynein Regulatory Complex (N-DRC) is a key regulator of ODA function, but the N-DRC component CCdc65 (Austin-Tse et al., 2013; Horani et al., 2013), and this protein displayed far faster kinetics than did the ODA subunits (Fig 5A). In fact, Ccdc65 kinetics were very similar to Enkur and Cfap52 (Fig. 5B), suggesting it may be continuously integrated as the axoneme assembles. Another example is the ODA docking complex protein, Ttc25 (Wallmeier et al., 2016). This protein is essential for the docking of ODAs to the axoneme, but it displayed far faster deployment kinetics than did either ODA subunit examined (Fig. 5C). This result suggests that the docking complex is deployed to axonemes in advance of the transport of the ODAs themselves. These results were confirmed by dual-labeling experiments (Fig. 5D-F).

**Figure 5:**
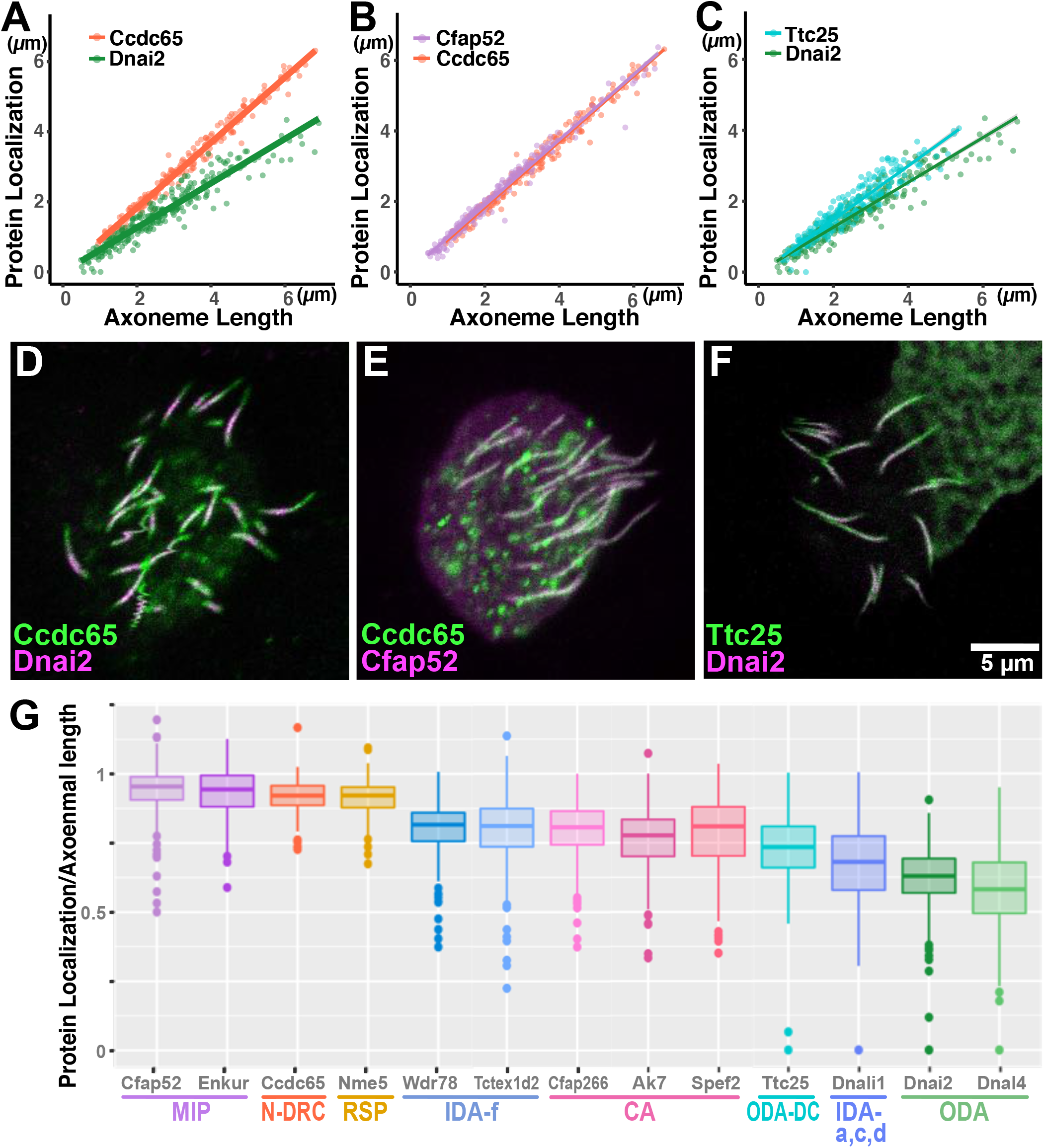
A-C) Additional elements of ciliary beating machineries display unexpected kinetics, and dual labeling confirms these relative rates (D-F). G) Summary of ciliary beating machinery deployment kinetics. Graph shows the mean ratio of decorated axoneme length versus total axoneme length for each protein.

## Discussion

The development of MCCs begins with their specification by an evolutionarily conserved transcriptional regulatory network and ultimately ends with the assembly and polarization of beating cilia (Brooks and Wallingford, 2014; Mahjoub et al., 2022). Here, we report the relative rates of axonemal deployment of diverse machines required for normal ciliary beating in growing MCC cilia in *Xenopus* (Fig. 5G). By combining quantification of axonemal deployment rates with two-color dual labeling of pairs of axonemal proteins, we find a consistent order to the deployment of these machineries. MIPs and radial spokes are deployed at the same rate as overall axoneme growth, suggesting they may be continuously integrated as the doublets are assembled. The central apparatus and “f” subtype of IDAs follow slightly behind the MIPs and radial spokes, with IDA(a,c,d) lagging slightly behind. The last elements to be deployed to developing MCC axonemes are the ODAs.

These results raise interesting questions about the relationships between beating machineries and IFT, which transports a wide array of ciliary cargoes and essential for ciliogenesis (Lechtreck, 2015). For example, tubulin for the outer doublet microtubules is a known cargo of IFT (e.g. (Bhogaraju et al., 2013; Craft et al., 2015; Marshall and Rosenbaum, 2001)), but recent work suggests that diffusion transports the majority of tubulin (Craft Van De Weghe et al., 2020). It will be interesting, then, to determine if IFT or diffusion accounts for the rapid deployment of MIPs into axonemes. Similarly, radial spokes are known cargoes of IFT (Lechtreck et al., 2022; Qin et al., 2004), yet they deploy at rates similar to that of the growing axonemes, raising the possibility that -like tubulin-they may also undergo substantial diffusive movement during axoneme growth.

Finally, the distinct rates observed for IDA and ODA deployment add yet another piece to the complex puzzle surrounding the regulation of these motors (Desai et al., 2018; Qiu and Roy, 2022). Dynein arms generally are thought to be transported by IFT (Hou et al., 2007; Pazour et al., 1998; Qin et al., 2004; Viswanadha et al., 2014), and our data suggest not only that ODAs and IDAs display widely divergent rates of axonemal deployment, but also that even different IDA sub-types display subtly distinct rates. These results in vertebrate MCCs is consistent with the findings in *Chlamydomonas* that while IDA deployment via the adaptor Ida3 is a length-dependent process (Hunter et al., 2018; Viswanadha et al., 2014), ODA deployment via the Oda16 adaptor is not (Ahmed et al., 2008; Hou and Witman, 2017; Taschner et al., 2017). It will be important and interesting to ask if other types of motile cilia in vertebrates such as mammalian airway or brain MCCs or nodal cilia which lack the central pair display similar patterns to those reported here.

## Acknowledgements

Thea authors thank Sarah Ortiz for initial characterization of the Dnali1 fusion protein. This work was supported by the NHLBI (R01HL117164).

## Materials and Methods

### *Xenopus* embryo manipulations

*Xenopus* embryo manipulations were carried out using standard protocols. Briefly, female adult *Xenopus* were induced to ovulate by injection of hCG (human chorionic gonadotropin). *In vitro* fertilization was carried out by homogenizing a small fraction of a testis in 1X Marc’s Modified Ringer’s (MMR). Embryos were dejellied in 1/3x MMR with 2.5 %(w/v) cysteine (pH7.8). Embryos were microinjected with mRNA in 2 % Ficoll (w/v) in 1/3x MMR and injected embryos were washed with 1/3x MMR after 30min.

### Plasmids for microinjections

*Xenopus* gene sequences were obtained from Xenbase (www.xenbase.org) and open reading frames (ORF) of genes were amplified from the *Xenopus* cDNA library by polymerase chain reaction (PCR), and then are inserted into a pCS10R vector containing a fluorescence tag. These constructs were linearized, and the capped mRNAs were synthesized using mMESSAGE mMACHINE SP6 transcription kit (ThermoFisher Scientific). 40∼80 pg of each mRNA was injected into two ventral blastomeres.

### Imaging and image analysis

*Xenopus* embryos were mounted between cover glass and submerged in 1/3x MMR at tadpole stages, and then were imaged immediately. Live images were captured with a Zeiss LSM700 laser scanning confocal microscope using a plan-apochromat 63X/1.4 NA oil objective lens (Zeiss) or with Nikon eclipse Ti confocal microscope with a 63×/1.4 oil immersion objective. Data were analyzed in Fiji and were visualized using RStudio.

